# Myogenic Vasoconstriction Requires Canonical G_q/11_ Signaling of the Angiotensin II Type 1a Receptor in the Murine Vasculature

**DOI:** 10.1101/2020.09.09.289280

**Authors:** Yingqiu Cui, Mario Kassmann, Sophie Nickel, Chenglin Zhang, Natalia Alenina, Yoland Marie Anistan, Johanna Schleifenbaum, Michael Bader, Donald G. Welsh, Yu Huang, Maik Gollasch

**Affiliations:** Charité - Universitätsmedizin Berlin, Experimental and Clinical Research Center (ECRC), a joint cooperation between the Charité Medical Faculty and the Max Delbrück Center for Molecular Medicine (MDC), Lindenberger Weg 80, 13125 Berlin, Germany; Heart and Vascular Institute and School of Biomedical Sciences, Chinese University of Hong Kong, Hong Kong, China; Max Delbrück Center for Molecular Medicine, Berlin, Germany; DZHK (German Center for Cardiovascular Research), Partner Site Berlin, Berlin, Germany; Charité - Universitätsmedizin Berlin, Germany; Institute for Biology, University of Lübeck, Lübeck, Germany; Robarts, Research Institute, Department of Physiology and Pharmacology, Western University, London, ON, Canada; Charité - Universitätsmedizin Berlin, Medical Clinic for Nephrology and Internal Intensive Care, Campus Virchow, 13353 Berlin, Germany; Department of Internal Medicine and Geriatrics, University Medicine Greifswald, Greifswald, Germany

**Keywords:** Angiotensin II type 1a receptor, biased ligands, myogenic vasoconstriction, arterial smooth muscle

## Abstract

**Background:** The myogenic response is an inherent vasoconstrictive property of resistance arteries to keep blood flow constant in response to increases in intravascular pressure. Angiotensin II (Ang II) type 1 receptors (AT1R) are broadly distributed, mechanoactivated receptors, which have been proposed to transduce myogenic vasoconstriction. However, the AT1R subtype(s) involved and their downstream G protein- and β-arrestin-mediated signaling pathways are still elusive.

**Objective:** To characterize the function of AT1aR and AT1bR in the regulation of the myogenic response of resistance size arteries and possible downstream signaling cascades mediated by G_q/11_ and/or β-arrestins.

**Methods:** We used *Agtr1a*^-/-^, *Agtr1b*^-/-^ and tamoxifen-inducible smooth muscle-specific AT1aR knockout mice (*SM-Agtr1a* mice). FR900359, [Sar1, Ile4, Ile8] Ang II (SII) and TRV120055 were used as selective G_q/11_ protein inhibitor and biased agonists to activate non-canonical β-arrestin and canonical G_q/11_ signaling of the AT1R, respectively.

**Results:** Myogenic and Ang II-induced vasoconstrictions were diminished in the perfused renal vasculature of *Agtr1a*^-/-^ and *SM-Agtr1a* mice. Similar results were observed in isolated pressurized mesenteric and cerebral arteries. Myogenic tone and Ang II-induced vasoconstrictions were normal in arteries from *Agtr1b*^-/-^ mice. The G_q/11_ blocker FR900359 decreased myogenic tone and Ang II vasoconstrictions while selective biased targeting of AT1R β-arrestin signaling pathways had no effects.

**Conclusion:** The present study demonstrates that myogenic arterial constriction requires G_q/11_-dependent signaling pathways of mechanoactivated AT1aR but not G protein-independent, noncanonical alternative signaling pathways in the murine mesenteric, cerebral and renal circulation.

## 1. Introduction

Myogenic vasoconstriction reflects the inherent ability of resistance arteries to adapt their diameter in response to alterations of intraluminal pressure. This response was first described by William Bayliss (2) and it reflects changes to the contractile state of vascular smooth muscle. Increases in transmural pressure cause vasoconstriction whereas decreases produce the opposing effect; this prototype of autoregulation has been observed in various microvascular arterial beds (8) and it is responsible for maintaining constant blood flow during fluctuations in perfusion pressure. Many cardiovascular disorders are associated with dysfunctional arterial myogenic response and they include hypertension, chronic heart failure, ischemic stroke, diabetes mellitus (6) (14) (31) (41) (45). Despite the functional importance of the myogenic response, the molecular mechanisms responsible for sensing intraluminal pressure has yet to be fully clarified.

Myogenic vasoconstriction is mediated by pressure-dependent depolarization of vascular smooth muscle cells, an event that augments Ca^2+^ influx through voltage-dependent Ca_v_1.2 channels (39) (7) (9) (16) (17) (38). G_q/11_-coupled receptors (GPCRs) are thought to function as the upstream sensor of membrane stretch (37), with angiotensin II type 1a (AT_1a_R), and perhaps AT1bR receptors in concert with cysteinyl leukotriene 1 receptor (CysLT1R), playing a particularly important role in the mesenteric and renal circulation (3) (40) (46) (49). AT1Rs are known to couple primarily to classical G_q/11_ proteins to activate multiple downstream signals, including protein kinase (PKC), extracellular signal-regulated kinases (ERK1/2), Raf kinases, tyrosine kinases, receptor tyrosine kinases (EGFR, PDGF, insulin receptor) and reactive oxygen species (ROS) (1). The AT1R activation also stimulates G protein-independent signaling pathways such as β-arrestin-mediated mitogen-activated protein kinase (MAPK) activation and Src-JAK/STAT (1). Recently, it has been shown that the activation of intracellular signaling by mechanical stretch of the AT1R does not require the natural ligand angiotensin II (Ang II) (44) (55) (46) but requires the activation of the transducer β-arrestin (44). Interestingly, mechanical stretch appears to allosterically stabilize specific β-arrestin-biased active conformations of AT1R to promote noncanonical downstream signaling mediated exclusively by the multifunctional scaffold protein, β-arrestin (50). Whether this noncanonical β-arrestin effector pathway plays a role in myogenic and ligand-dependent vasoconstriction has yet to be ascertained.

This study explored the specific function of AT1R subtypes in the regulation of myogenic tone and whether downstream signaling pathways are dependent on canonical G_q/11_ and/or noncanonical alternative signaling pathways. In this regard, we generated mice with cell specific deletion of smooth muscle AT1a receptors (*SM-Agtr1a* mice) and studied the effects of biased GPCR agonists and G_q/11_ protein inhibition on tone development in three distinct vascular beds (renal, cerebral and mesenteric circulation). We found that the AT1aR coupled towards the canonical G_q/11_ signaling pathway is required for the myogenic response in all three vascular beds. Our data argue against involvement of noncanonical G protein-independent alternative signaling downstream of the AT1aR to cause myogenic vasoconstriction.

## 2. Materials and Methods

### 2.1 Mouse Model

We used the SMMHC-Cre-ER^T2^ transgenic mouse line expressing Cre recombinase in smooth muscle cells under control of the smooth muscle myosin heavy chain promoter (26) and a mouse line bearing a floxed allele of the *Agtr1a* gene (*Agtr1a*^flox^), encoding the major murine AT1 receptor isoform (AT1aR) (48) to generate SMMHC-Cre+*Agtr1a*^flox/flox^ (*SM-Agtr1a*^-/-^) mice (**Figure 1A**). Genotyping was performed by polymerase chain reaction (PCR) analysis of tail DNA as described previously (26). Amplification of the SMMHC-Cre gene was performed in a multiplex PCR with the primers TGA CCC CAT CTC TTC ACT CC (SMWT1), AAC TCC ACG ACC ACC TCA TC (SMWT2), and AGT CCC TCA CAT CCT CAG GTT (phCREAS1) (13). The following primers (5’-3’) were used to identify *Agtr1a*^flox^ alleles: forward GCT TTC TCT GTT ATG CAG TCT, reverse ATC AGC ACA TCC AGG AAT G. Adult (12-16 weeks) male mice were injected with tamoxifen (30 μg/mg body weight) on 5 consecutive days. Isolated arteries were usually obtained after 2 to 3 days after tamoxifen treatment. **Figure 1B** shows reduction of AT1aR expression in vascular smooth muscle cells of *SM-Agtr1a*^-/-^ arteries. We also studied adult (12-16 weeks) male mice with global AT1a receptor deficiency (*Agtr1a*^-/-^) (24) (46) (25), and with global AT1b receptor deficiency (*Agtr1b*^-/-^) (40). Age-matched male mice were used as controls in the experiments. Animal care followed American Physiological Society guidelines, and all protocols were approved by local authority (LAGeSo, Berlin, Germany) and the animal welfare officers of the Max Delbruck Center for Molecular Medicine. Mice were maintained in the Max Delbrück Center animal facility in individually ventilated cages (Tecniplast, Deutschland) under standardized conditions with an artificial 12-hour dark-light cycle, with free access to standard chow (0.25% sodium; SSNIFF Spezialitäten, Soest, Germany) and drinking water. Animals were randomly assigned to the experimental procedures.

**Figure 1:**
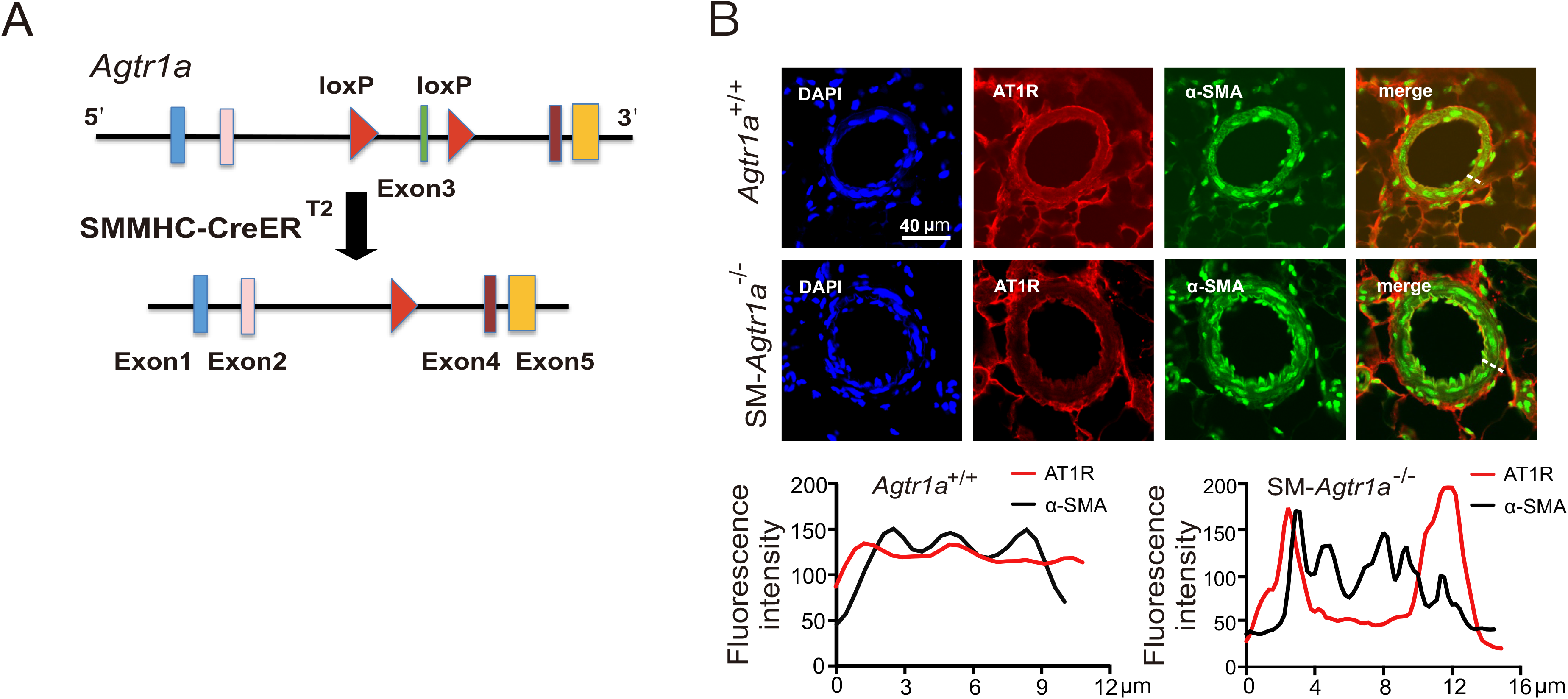
Conditional deletion of AT1a receptors in vascular smooth muscle cells of arteries. **A**: Schematic representation of the mouse allele containing loxP sequences, and the floxed allele after the action of Cre recombinase. **B:** Immunofluorescence staining results show that AT1R (red) is highly expressed in the mesenteric artery of *Agtr1a*^+/+^ mice. In *SM-Agtr1a*^-/-^ mouse mesenteric artery, the expression of AT1R is specifically reduced in smooth muscle cells. Scale bar: 40 μm.

### 2.2 Materials

Antibody to α-smooth muscle actin (α-SMA, #ab8211) was from Abcam (Cambridge, MA, USA). Anti-AT1R (#PA5-20812) and donkey anti-rabbit IgG (H+L) secondary antibody (A10040) were purchased from Thermo Fisher Scientific (Waltham, MA, USA). 4’,6-diamidino-2-phenylindole (DAPI, #D9542) was purchased from Sigma-Aldrich Co. (St. Louis, MO, USA). Ang II (#A9525), SII (#sc-391239A) and tamoxifen (#H7904) were from Sigma-Aldrich Co (82024 Taufkirchen, Germany). TRV120055 (#JT-71995) and TRV120056 (#JT-71996) were from Synpeptide Co., Ltd (Shanghai, China).

### 2.3 Mesenteric and cerebral arteries

After mice were killed, the mesenteric bed and brain were removed and placed into cold (4°C), gassed (95% O_2_-5% CO_2_) physiological saline solution (PSS) of the following composition (mmol/L): 119 NaCl, 4.7 KCl, 25 NaHCO_3_, 1.2 KH_2_PO_4_, 1.6 CaCl_2_, 1.2 MgSO_4_, 0.03 EDTA, and 11.1 glucose. Third or fourth order mesenteric and middle cerebral arteries or posterior cerebral arteries were dissected and cleaned of adventitial connective tissue (46) (18) (10) (11).

### 2.4 Pressure myography

Vessel myography was performed as previously described (26) (37) (46) (10). Mesenteric or cerebral arteries were mounted on glass cannula and superfused continuously with PSS (95% O_2_-5% CO_2_; pH, 7.4; 37°C). The vessels were stepwise pressurized to 20, 40, 60, 80, or 100 mmHg using a pressure servo control system (Living System Instrumentation, Burlington, VT). We measured the inner diameter of the vessels with a video microscope (Nikon Diaphot, Düsseldorf, Germany) connected to a personal computer for data acquisition and analysis (HaSoTec, Rostock, Germany) (18) (19) (46) (11) (10). Arteries were equilibrated for 45 to 60 minutes before starting experiments. A 60-mmol/L KCl challenge was performed before any other intervention.

### 2.5 Analysis of myogenic tone in isolated perfused kidneys

Isolated kidneys were perfused in an organ chamber using a peristaltic pump at constant flow (0.3-1.9 ml/min) of oxygenated (95% O_2_ and 5% CO_2_) PSS (46). Drugs (Ang II or biased agonists) were added to the perfusate. Perfusion pressure was measured by a pressure transducer after an equilibration period of 60-90 min. Data were recorded and analyzed by a Powerlab acquisition system (AD Instruments, Colorado Springs). Ang II-induced pressor effects were normalized to the maximal pressor effect induced by KCl (60 mmol/L) (37) (46) (18).

### 2.6 Immunofluorescence

*Agtr1a*^+/+^ and *SM-Agtr1a*^-/-^ mice mesenteric arteries were dissected and further fixed in 4% formaldehyde and embedded in Tissue-Tek O.C.T. compound to be frozen in liquid nitrogen. Tissues were then sectioned and permeabilized in 1% Triton X-100 in PBS. Sections were stained with the primary antibody overnight at 4°C. After washing with PBS for 3×5 min, the secondary antibody and DAPI were applied for 2 hours at room temperature. Fluorescence images were captured by use of Olympus FV1000 confocal microscopy and images were analyzed by ImageJ analysis software.

### 2.7 Statistics

Data are presented as means ± SEM. Statistically significant differences in mean values were determined by Student’s unpaired t test or one-way analysis of variance (ANOVA). P values < 0.05 were considered statistically significant.

## 3. Results

### 3.1 AT1aR is essential for pressure-induced response in the renal circulation

We evaluated myogenic tone in mouse renal circulation, a highly myogenic bed regulating blood flow to the kidneys and consequently sodium excretion and systemic blood pressure. Renal vascular resistance of isolated perfused kidneys was determined by measuring perfusion pressure at fixed levels of flow. The perfusion pressure increased with flow rate in kidneys of wild-type *Agtr1a*^+/+^ mice, reaching a value of about 160 mmHg at a flow rate of 1.9 ml/min (**Figure 2A**). Kidneys from *Agtr1a*^-/-^ mice developed significantly less pressure at the same flow rate (**Figure 2B, E**). 60 mmol/L KCl-induced increases in perfusion pressure were normal in *Agtr1a*^-/-^ kidneys (**Figure 2 F**). At a flow rate of 1.9 ml/min, pressure in *Agtr1a*^-/-^ kidneys was ~100 mmHg lower than in *Agtr1a*^+/+^ kidneys. Angiotensin II (Ang II, 10 nmol/L) increased perfusion pressure by ~80 mmHg in kidneys of *Agtr1a*^+/+^ mice, but had no effect in kidneys of *Agtr1a*^-/-^ mice (**Figure 2C**); this is indicative of AT1aRs mediating Ang II-dependent vasoconstriction. Removal of external Ca^2+^ nearly abolished flow-induced myogenic constriction in perfused kidneys of *Agtr1a*^+/+^ mice, but had nearly no effect in kidneys of *Agtr1a*^-/-^ mice (**Figure 2D**), indicating AT1aRs mediate also myogenic constriction of mouse renal arterioles. Of note, there was no difference in myogenic tone and Ang II vasoconstrictions between *Agtr1b*^-/-^ versus *Agtr1b*^+/+^ kidneys (**Figure 3**).

**Figure 2:**
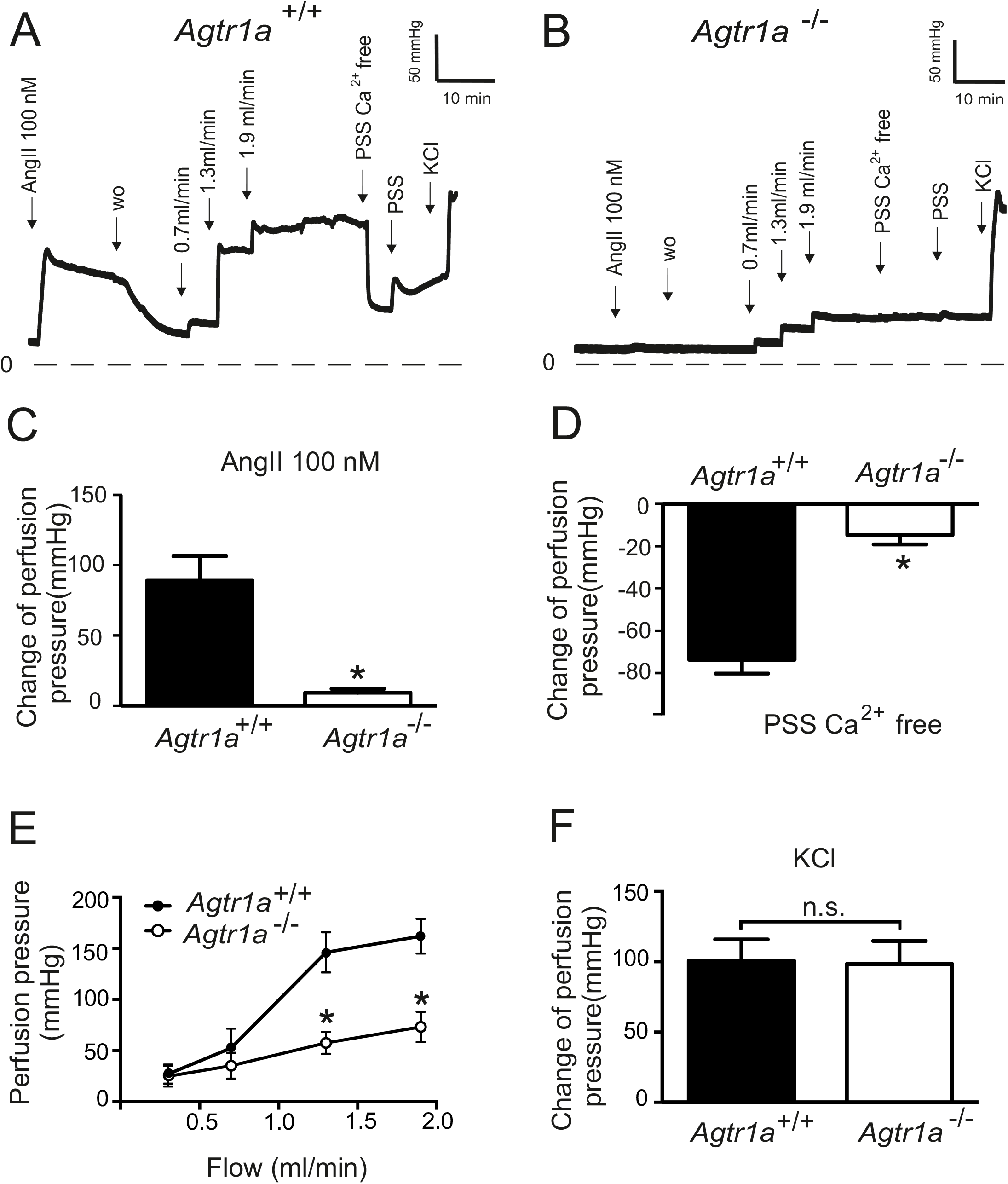
Vasoregulation in isolated perfused kidneys of *Agtr1a*^-/-^ mice. **A, B**: Original recordings of perfusion pressure in kidneys of *Agtr1a*^+/+^ (**A**) and *Agtr1a*^-/-^ mice (**B**). **C:** Increase in the perfusion pressure induced by 100 nM Ang II. **D:** Myogenic tone assessed by exposure to Ca^2+^ free PSS. **E**: Perfusion pressure at flow rates of 0.3 ml/min, 0.7 ml/min, 1.3 ml/min and 1.9 ml/min. **F**: Increase in the perfusion pressure induced by 60mM KCl. n=6 *Agtr1a*^+/+^ kidneys and n=7 *Agtr1a*^-/-^ kidneys for all panels. *p<0.05; n.s., not significant.

**Figure 3:**
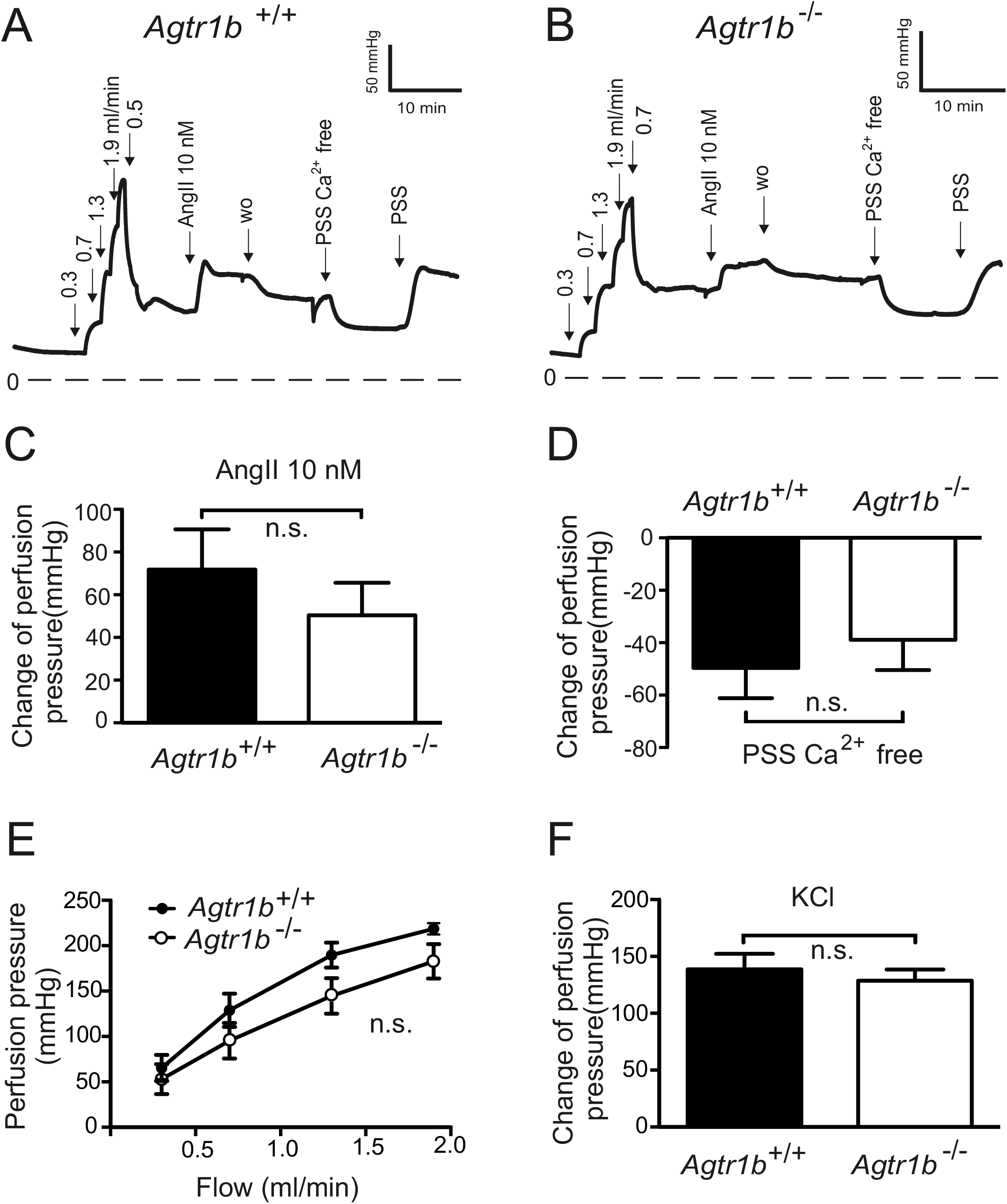
Vasoregulation in isolated perfused kidneys of *Agtr1b*^-/-^ mice. **A, B**: Original recordings of the perfusion pressure in kidneys of *Agtr1b*^+/+^ (**A**) and *Agtr1b*^-/-^ mice (**B**). **C:** Increase in perfusion pressure induced by 10 nM Ang II. **D:** Change of pressure assessed by exposure to Ca^2+^ free PSS. **E**: Perfusion pressure at flow rates of 0.3 ml/min, 0.7 ml/min, 1.3 ml/min and 1.9 ml/min. **F**: Increase in perfusion pressure induced by 60 mM KCl. n=6 *Agtr1b*^+/+^ kidneys and n=6 *Agtr1b*^-/-^ kidneys for all panels. w.o., wash-out; n.s., not significant.

Next we focused on kidneys from *SM-Agtr1a*^-/-^ mice (**Figure 4**). At a flow rate of 1.9 ml/min, pressure in *SM-Agtr1a*^-/-^ kidneys was ~90 mmHg lower than in *Agtr1a*^+/+^ kidneys (**Figure 4 A, B, E**). *SM-Agtr1a*^-/-^ kidneys showed largely reduced myogenic vasoconstriction as assessed by exposure of the kidneys to Ca^2+^ free PSS (**Figure 4B, D**), whereas wild-type kidneys showed strong myogenic vasoconstrictions (**Figure 4A, D**). 60 mmol/L KCl-induced increases in perfusion pressure were normal in *Agtr1a*^-/-^ kidneys (**Figure 4F**). Ang II (10 nmol/L) induced weaker increases in perfusion pressure in kidneys of *SM-Agtr1a*^-/-^ mice compared to controls (**Figure 4D**). Together, these results reveal a key role of AT1aR but not AT1bR in the flow-induced myogenic response of the mouse renal vasculature.

**Figure 4:**
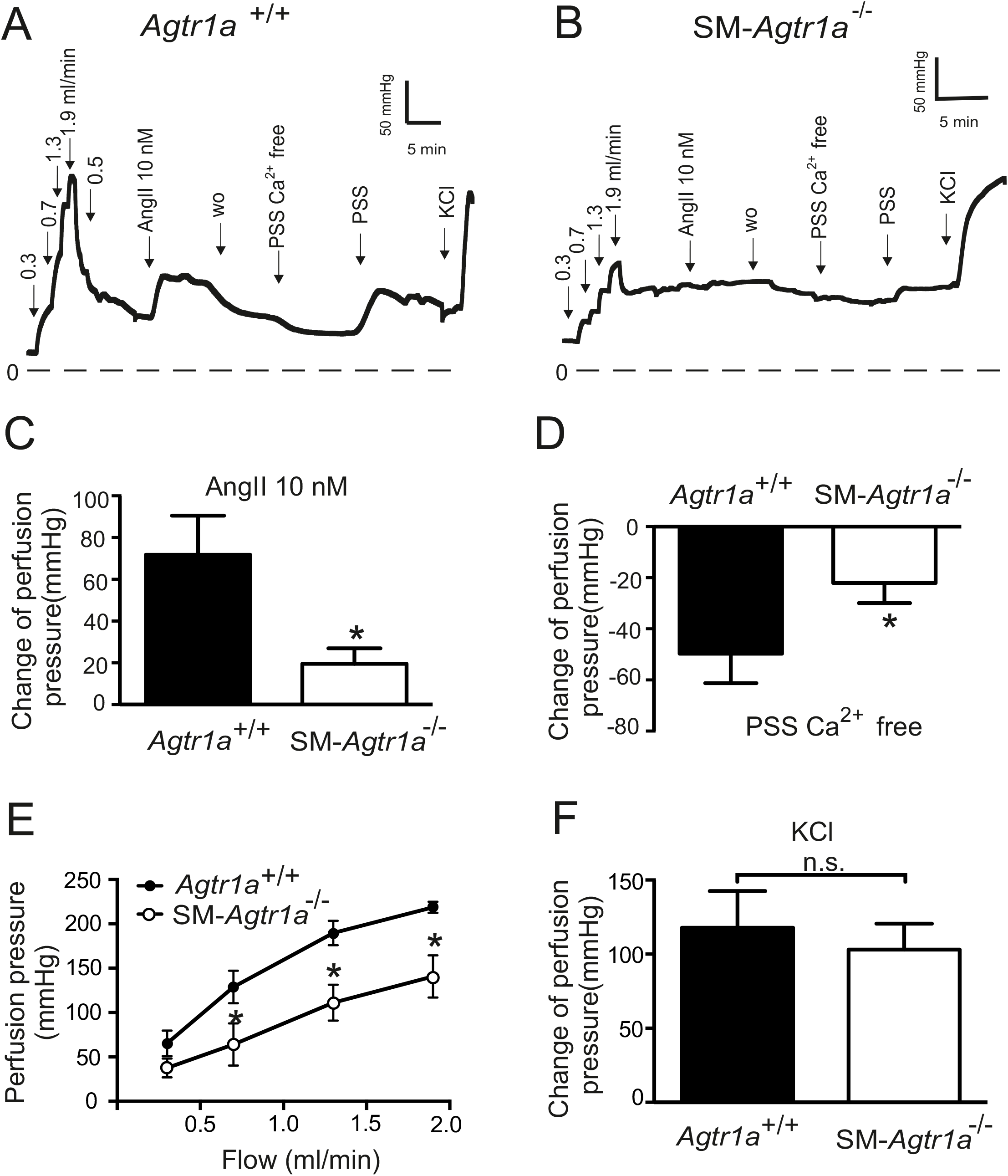
Vasoregulation in isolated perfused kidneys of *SM-Agtr1a*^-/-^ mice. **A, B**: Original recordings of the perfusion pressure in kidneys of *Agtr1a*^+/+^ (**A**) and *SM-Agtr1a*^-/-^ mice (**B**). **C:** Increase in perfusion pressure induced by 10 nM Ang II. **D:** Change of pressure assessed by exposure to Ca^2+^ free PSS. **E**: Perfusion pressure at flow rates of 0.3 ml/min, 0.7 ml/min, 1.3 ml/min, and 1.9 ml/min. **F**: Increase in perfusion pressure induced by 60 mM KCl. n=6 *Agtr1a*^+/+^ kidneys and n=6 *SM-Agtr1a*^-/-^ kidneys for all panels. *p<0.05; n.s., not significant.

### 3.2 AT1aR contribute to myogenic constriction in mesenteric arteries

We monitored myogenic constriction in resistance-sized mesenteric arteries using videomicroscopy. Mesenteric arteries were exposed to stepwise (20 mmHg) increases in intraluminal pressure (20-100 mmHg) in the presence and absence of external Ca^2+^ (1.6 mmol/L) to determine active and passive vessel diameters, respectively. **Figure 5** shows representative recordings of mesenteric arteries from *Agtr1a*^+/+^ mice (**Figure 5A**) and *SM-Agtr1a*^-/-^ mice (**Figure 5B**) and myogenic vasoconstriction was defined as the diameter difference in the presence and absence of external Ca^2+^ (1.6 mmol/L) at each pressure step (46). Increases in intraluminal pressure generated active tension that counteracted further dilation of the vessels at 60 to 80 mmHg in mesenteric arteries from *Agtr1a*^+/+^ mice, reaching peak constrictions of 50 μm at 80 to 100 mmHg (**Figure 5A**). In contrast, mesenteric arteries from *SM-Agtr1a*^-/-^ mice only produced ~35% of the constriction observed in wild-type arteries (**Figure 5B, C**). Ang II strongly constricted arteries from *Agtr1a*^+/+^ mice but had no effect on arteries from *SM-Agtr1a*^-/-^ mice (**Figure 5D**); the latter did constrict in response to 60 mmol/L KCl (**Figure 5E**). This study observed a marked reduction in AT1aR expression in the media of *SM-Agtr1a*^-/-^ mesenteric arteries compared to wild-type (**Figure 1B**), in keeping with this receptor mediating myogenic constriction in mesenteric arteries.

**Figure 5:**
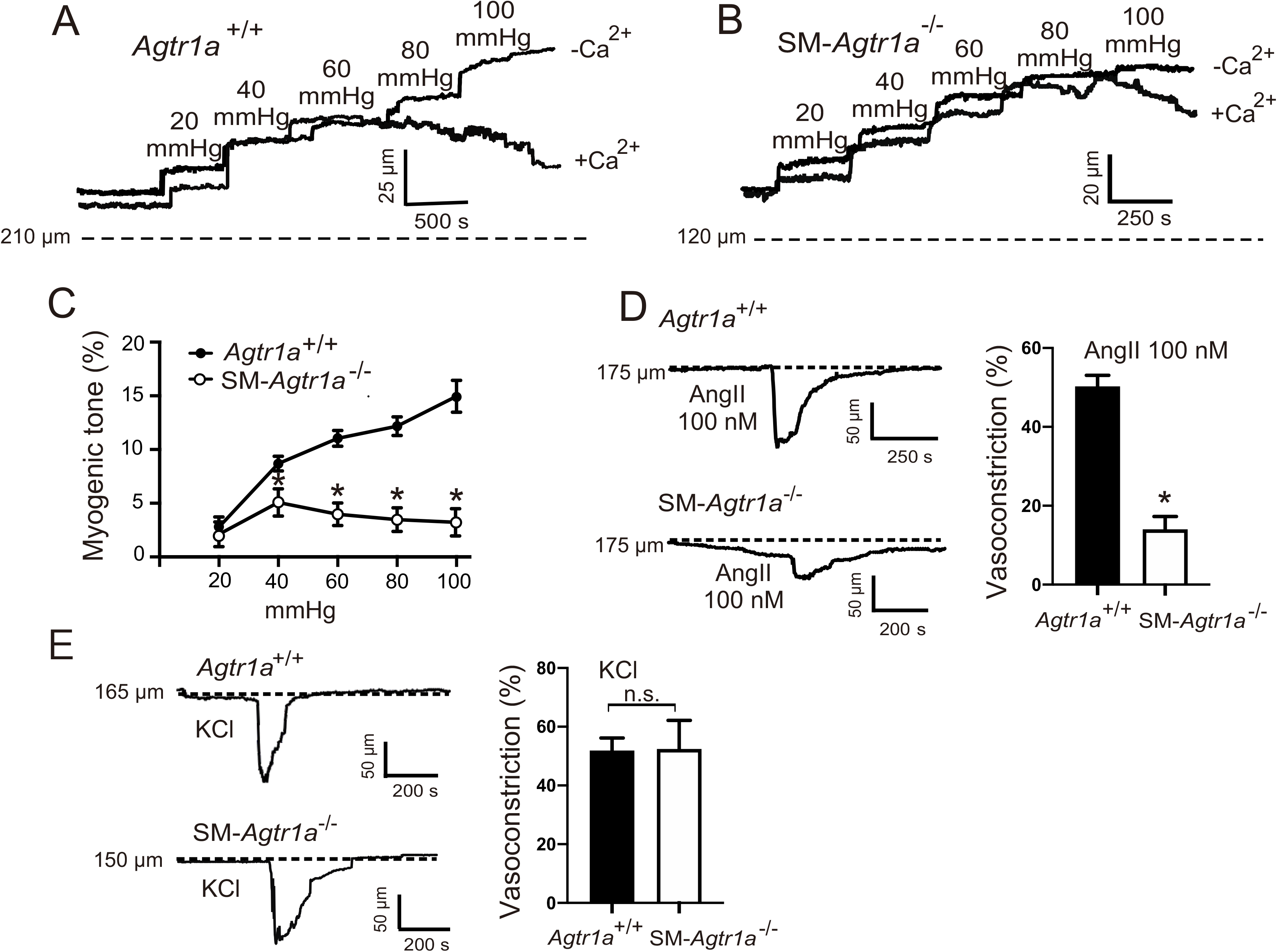
Myogenic tone in mesenteric arteries. **A, B**: Representative recordings of MA diameter during a series of pressure steps from 20 to 100 mmHg in 20 mmHg increments in control conditions (+Ca^2+^) and in Ca^2+^ free solution (-Ca^2+^). Arteries were isolated from *Agtr1a*^+/+^ **(A)** and *SM-Agtr1a*^-/-^ mice **(B)**. Note the increase in active constriction over the entire pressure range from 60 to 100 mmHg in vessels from Agtr1a^+/+^, but not from *SM-Agtr1a*^-/-^ mice. Vasodilation in Ca^2+^-free solution was observed in Agtr1a^+/+^ but not in *SM-Agtr1a*^-/-^ arteries (P<0.05). **C**: Myogenic tone (at 80 mmHg) expressed as dilation of vessels induced by external Ca^2+^ free solution (0 Ca/EGTA; n=6). **D** to **G**: Response to angiotensin II (Ang II; **D, E**) and 60 mM KCl (**F, G**) in MA of *Agtr1a*^+/+^ and *SM-Agtr1a*^-/-^ mice. MAs were pressurized to 60 mmHg. Responses are expressed as relative changes in vessel inner diameter. *Agtr1a^+/+^,* n=5 vessels and *SM-Agtr1a^-/-^,* n=4 vessels for each group. *p<0.05.

### 3.3 AT1aR contribute to myogenic constriction in cerebral arteries

Next, we studied the function of AT1aRs in cerebral arteries. Vessels were equilibrated at 15 mmHg (30 min) and following an assessment of KCl-induced constriction, arteries were pressurized to 80 mmHg (**Figure 6A**). Ang II constrictions and myogenic constriction was significantly decreased in *SM-Agtr1a*^-/-^ arteries compared to wild-type (**Figure 6A, B, C, D**). Both wild-type and *SM-Agtr1a*^-/-^ arteries produced similar constrictions when exposed to 60 mmol/L KCl (**Figure 6E**). The results demonstrate a key role of AT1aR in the myogenic response of mouse cerebral arteries.

**Figure 6:**
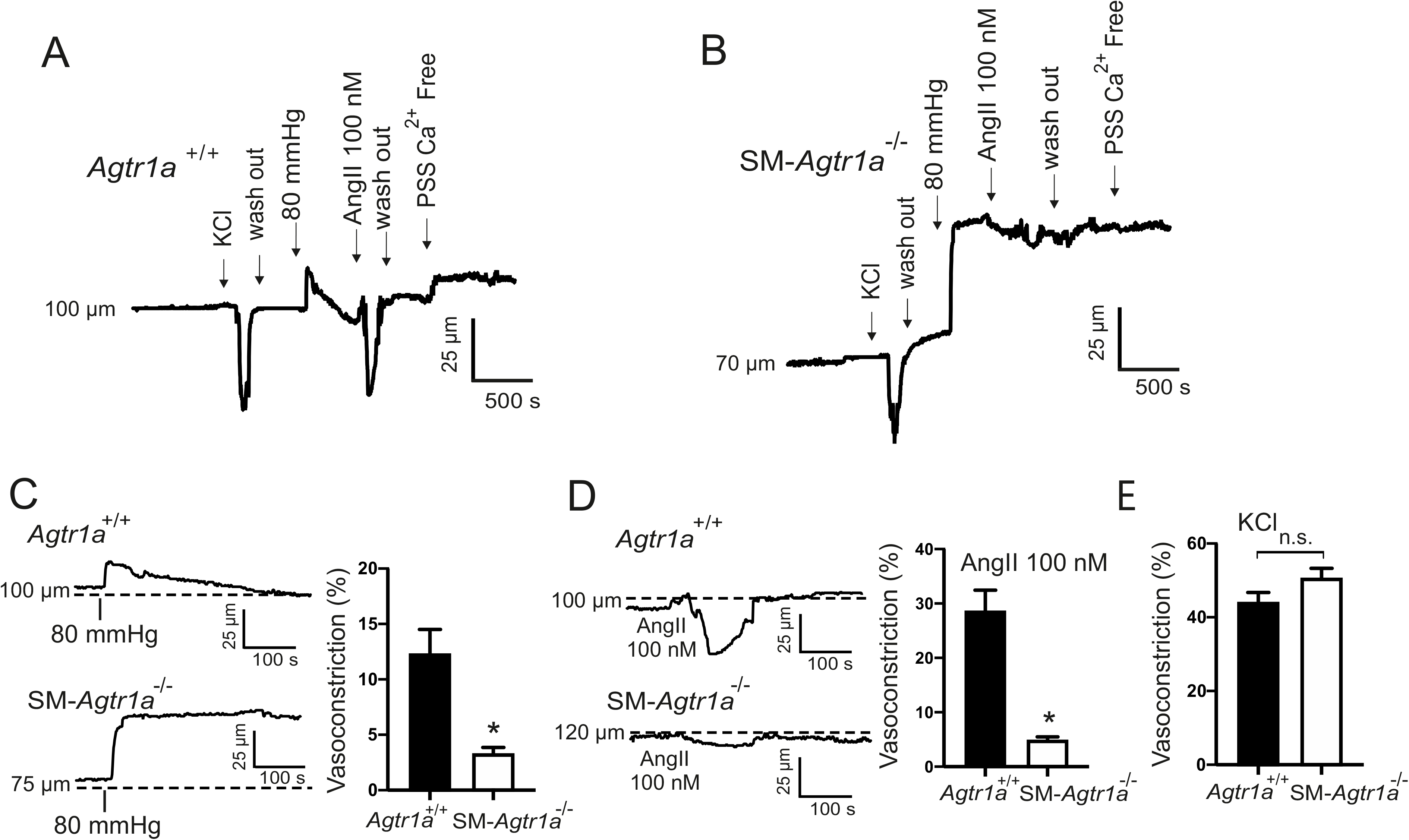
Myogenic tone in cerebral arteries. **A, B**: Representative recordings of middle/posterior cerebral arteries diameter at the pressure of 80 mmHg in control conditions (WT), Ang II 100 nmol/L, and in Ca^2+^ free solution. **C**: Myogenic tone (at 80 mmHg) expressed as dilation of vessels induced by external Ca^2+^ free solution. **D, E**: Response to Ang II **(D)** and 60 mM KCl **(E)** in middle/posterior cerebral arteries of *Agtr1a*^+/+^ and *SM-Agtr1a*^-/-^ mice. *Agtr1a^+/+^,* n=6 vessels and *SM-Agtr1a^-/-^,* n=6 vessels for each group. * p<0.05.

### 3.4 G_q/11_ protein dependent signaling pathway is responsible for myogenic tone

To explore the role of G_q/11_ and β-arrestin signaling pathways downstream of AT1R, we used the biased agonists TRV120055 and SII to activate G_q/11_ and β-arrestin signaling pathways, respectively (29) (34) (51). We found that TRV120055 increased vascular tone in mesenteric arteries (**Figure 7A, B)**, whereas SII had no effect (**Figure 7C, D**). Similarly, TRV120055 and TRV120056 (another biased G_q/11_ coupled AT1R agonist) enhanced dose-dependent perfusion pressure in isolated kidneys (**Figure 8A, C**), whereas SII had no effect (**Figure 8B, C**). The removal of external Ca^2+^ abolished agonist-induced vasoconstrictions in perfused kidneys (**Figure 8D**), indicating AT1aRs mediate vasoconstriction *via* canonical G_q/11_ but not noncanonical β-arrestin pathways. To confirm the results, we next examined the effects of FR900359, a selective G_q/11_-protein inhibitor (23) (47) (35). FR900359 abolished both myogenic and Ang II-dependent constrictions in renal arterioles (**Figure 9**) and mesenteric arteries (**Figure 7E, F**). These results indicate that myogenic vasoconstriction is mediated through the mechanosensitive AT1aR and the canonical G_q/11_ signaling pathway.

**Figure 7:**
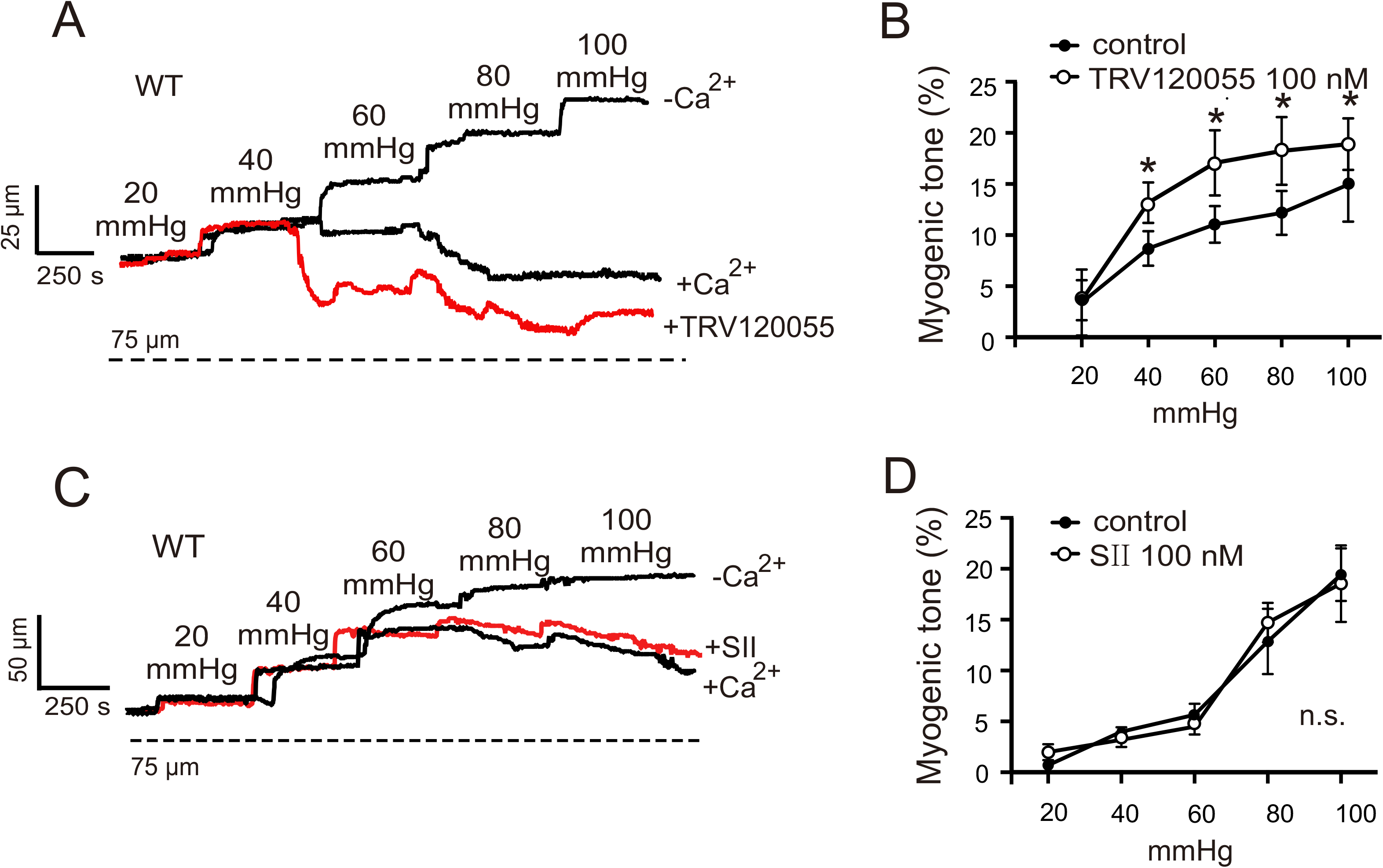
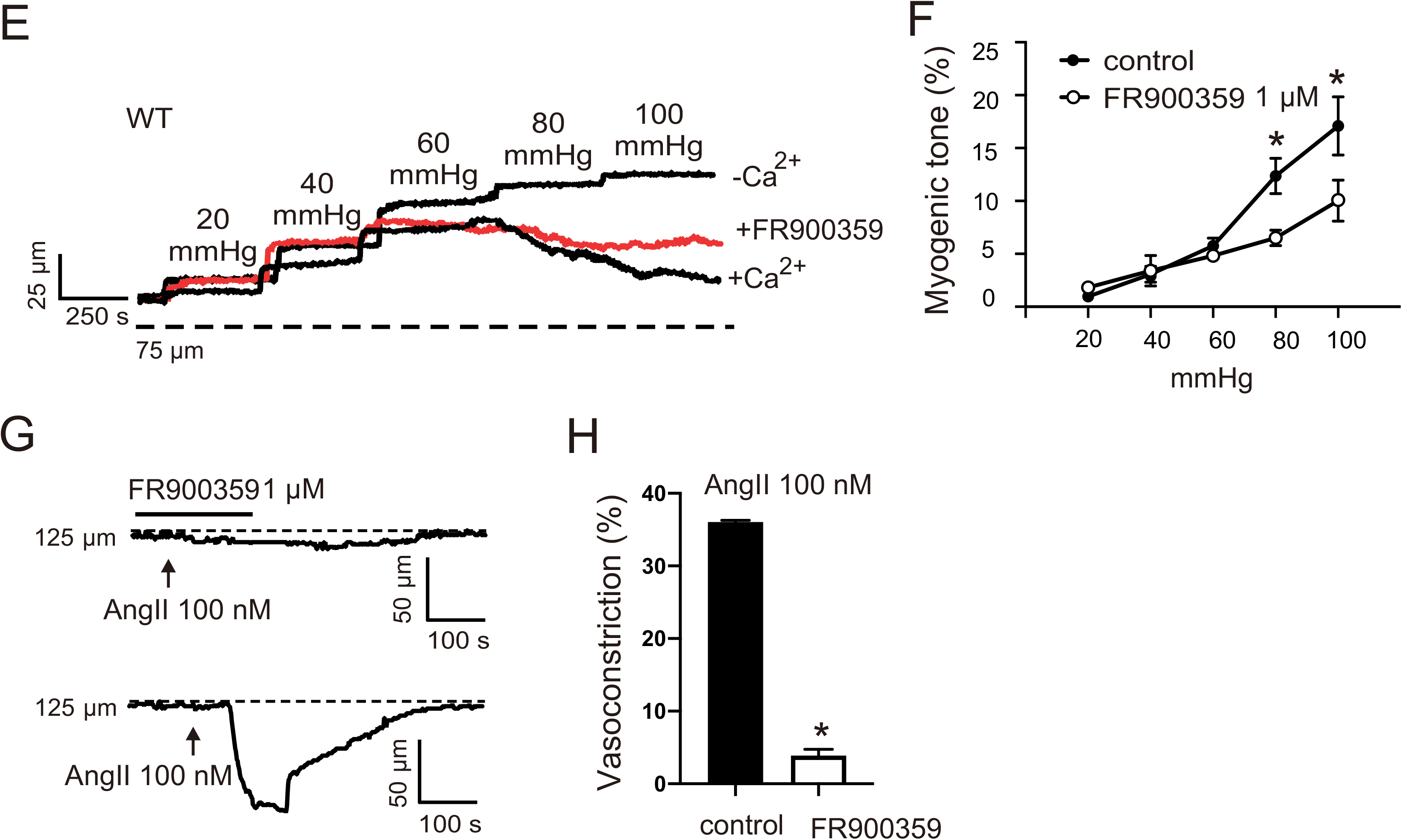
Enhancement of the vascular tone by TRV120055. **A, C, and E** Representative recordings of mesenteric artery diameter during a series of pressure steps from 20 to 100 mmHg in 20 mmHg increments in control conditions (+Ca^2+^), TRV120055 100 nmol/L **(A)**, SII 100 nmol/L **(C)**, FR120055 1 μmol/L **(E)** and in Ca^2+^-free solution. **B, D and F**: Average myogenic constriction of mesenteric arteries in drug-free physiological salt solution (PSS) and in PSS containing 100 nmol/L TRV120055 **(B)**, 100 nmol/L SII **(D)**, and 1 μmol/L FR120055 **(F)** (n=6, 4 and 4, respectively). **G and H:** Response to Ang II in MA in drug-free PSS and PSS in presence of FR120055 at 80 mmHg (n=6 each). *p<0.05; n.s., not significant.

**Figure 8:**
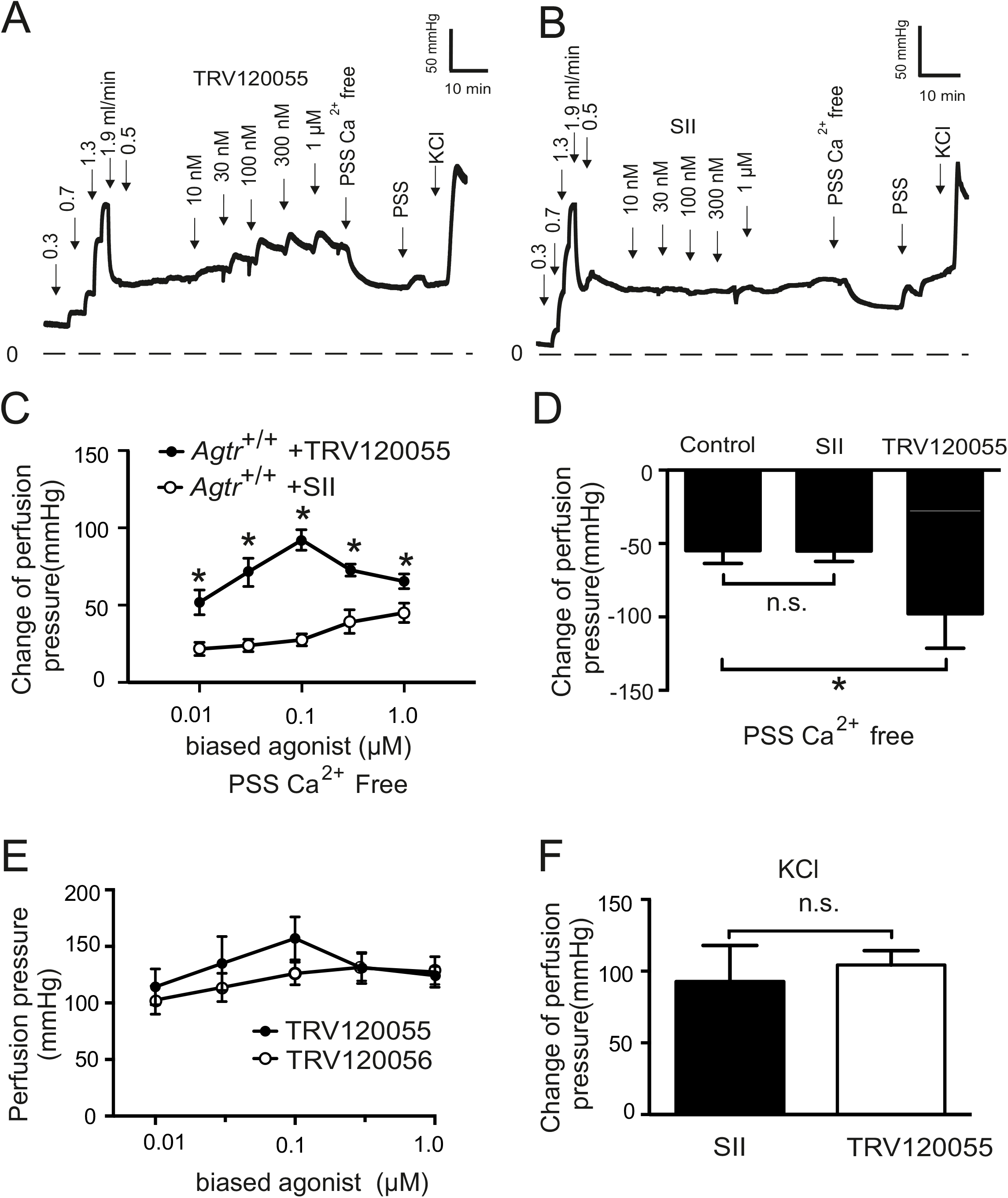
Function of biased AT1R agonists to vasoregulation in isolated perfused kidneys from *Agtr1a*^+/+^ mice. **A, B**: Original recordings of perfusion pressure in response to various flow rates (in ml/min), TRV120055 (**A**) or Sar-Ile II (**B**), Ca^2+^ free perfusion solution (PSS Ca^2+^ free) and re-exposure of the kidneys to PSS. **C:** Increase in perfusion pressure induced by TRV120055 and Sar-Ile II in various concentrations (10 nM to1 μM). **D:** Change of perfusion pressure assessed by exposure of the kidneys to Ca^2+^ free PSS at the presence of TRV120055 or Sar-Ile II at the concentration of 100 nM. **E**: Dose-response relationships for TRV120055 and TRV120056. **F**: Increase in perfusion pressure induced by 60 mM KCl. TRV120055, TRV120056, Sar-Ile II. n=6 kidneys in each group; n=6 kidneys in the control group. *p<0.05; n.s., not significant; Control, *Agtr*^+/+^ without biased ligand.

**Figure 9:**
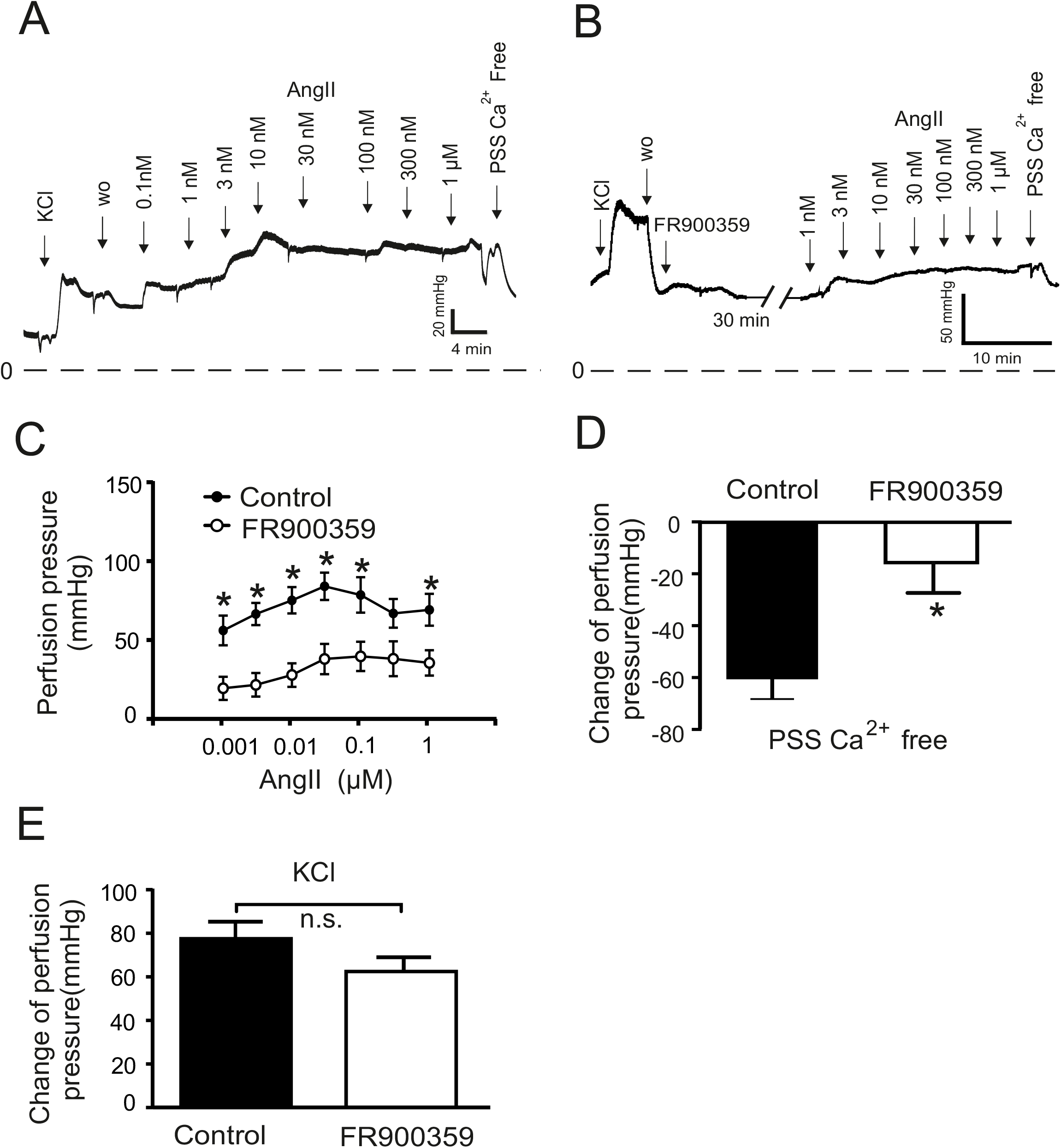
Vasoregulation in isolated perfused kidneys of *Agtr1a*^+/+^ mice pretreated with 300 nM G_q/11_ blocker FR900359. **A**: Original recordings of perfusion pressure in kidneys of *Agtr*^+/+^ mice in response to various concentrations of Angiotensin II (Ang II) **B**: same as A but pretreated with 300 nM FR900359 for 30 minutes. **C:** Increases in perfusion pressure induced by Ang II (1 nM to 1 μM). **D:** Myogenic tone assessed by exposure of the kidneys to Ca^2+^-free PSS. **E**: Increase in perfusion pressure induced by 60 mM KCl. n=5 *Agtr*^+/+^ kidneys and n=6 *Agtr1a*^+/+^ kidneys pretreated with FR900359 for all panels. *p<0.05; w.o., wash-out; n.s., not significant.

## 4. Discussion

The study found that the canonical G_q/11_ signaling of mechanoactivated AT1aR is responsible for myogenic vasoconstriction in mesenteric, renal arteries and cerebral arteries. We observed a loss of myogenic autoregulation in the renal circulation of *Agtrla*^-/-^ mice, an effect which was normal in *Agtrlb*^-/-^ mice. Similarly, we found that myogenic tone was strongly reduced in two other myogenic arteries (mesentery and cerebral) from smooth muscle specific AT1aR-deficient *(SM-Agtr1a^-/-^)* mice compared to wild-type. Using the pharmacological G_q/11_ inhibitor FR900359 and several GPCR biased agonists, we showed that AT1Rs cause vasoconstriction *via* canonical G_q/11_ signaling but not alternative G protein signaling downstream of the AT1R.

### ATIaRs are primary mechanosensors in intact arteries

Multiple GPCRs have been proposed to act as mechanosensors to regulate myogenic tone in resistance arteries. While stretch induces activation of purinergic P2Y6 UDP receptors, thromboxane A2 (TP) receptors and sphingosine-1-phosphate (S1P) receptors in certain vascular beds (27) (28) (30), the AT1R remains one of the best characterized mechanosensor in the vasculature (49) (55). Humans express a single type of AT1R, whereas two isoforms (AT1aR and AT1bR) are present in rodents (36) (53). Using *Agtrla*^-/-^ mice and inverse AT1R agonist, our previous data suggested that ligand-independent AT1aR activation is required for myogenic response in resistance mesenteric arteries and renal arterioles (46). However, two recent studies reported that myogenic tone was diminished in *Agtrlb*^-/-^ mesenteric and cerebral arteries, which implies a possible role of AT1bRs in mechanosensation (42) (3). In contrast, we found that myogenic tone was normal in *Agtrlb*^-/-^ perfused kidneys, which argues against a role of AT1bR in myogenic constriction in the renal circulation. This data was, however, obtained in global mutant mice, which often display compensatory mechanisms for the lack of AT1Rs. Moreover, AT1aR and AT1bR are expressed at similar levels in cerebral parenchymal arterioles and genetic knockout of AT1aR (but not AT1bR) blunted the ability of these vessels to generate myogenic tone (52). The latter effect is opposite to cerebral arteries where genetic knockout of AT1bR blunted the ability to develop myogenic tone (42). To overcome these potential limitations, we generated tamoxifen-inducible *SM-Agtr1a* (SMMHC-Cre+*Agtr1a*^flox/flox^) mice for careful phenotypic investigation. We found that myogenic constriction was impaired in cerebral, mesenteric and renal arteries isolated from smooth muscle AT1aR-deficient mice. The data provide firm evidence that AT1aRs play a key role as mechanosensors mediating myogenic constriction in the murine vasculature.

### AT1aRs downstream signaling to cause vasoconstriction

We next explored downstream signaling pathways mediated by G_q/11_ and/or β arrestins of the AT1R in the vascular response. In cell culture, osmotic cell stretch has been found to increase the binding affinity and potency of the β-arrestin-biased agonist TRV120023 with no effect on the balanced agonist Ang II through AT1R to induce a conformation change of β-arrestin 2, similar to that induced by β-arrestin-biased agonists (50). Similarly, hypo-osmotic stretch induced β-arrestin-biased signaling of AT1Rs in the absence of G protein activation (44). We failed to observe β-arrestin mediated enhancement of myogenic vasoconstriction with the β-arrestin biased agonist SII in intact arteries (mesenteric and renal arteries: **Figure 10**). The discrepancy might be caused by differences between the hypo-osmotic cell swelling and tensile stretch on the smooth muscle cell layer in intact arteries to cause mechanoactivation of AT1aRs *in situ*. GPCRs biased mechanisms have been described between two different G proteins, between β-arrestin-1 and 2, and between different states of the same receptor bound to different ligands (12) (21) (54). However, the majority of well described GPCRs biased ligand examples refers to selective G protein signaling versus β-arrestin-mediated signaling (20) (33) (43) (51). AT1aR is one of the best characterized GPCR enabling biased receptor signaling. It can be activated in either a canonical G protein-dependent signaling mode (5) (37) or noncanonical β-arrestin-mediated signaling mode (44) (50). In line, we found that the natural biased agonist Ang II was able to increase G protein signaling of mechanoactivated AT1R receptors to enhance the vasoconstrictor response.

**Figure 10:**
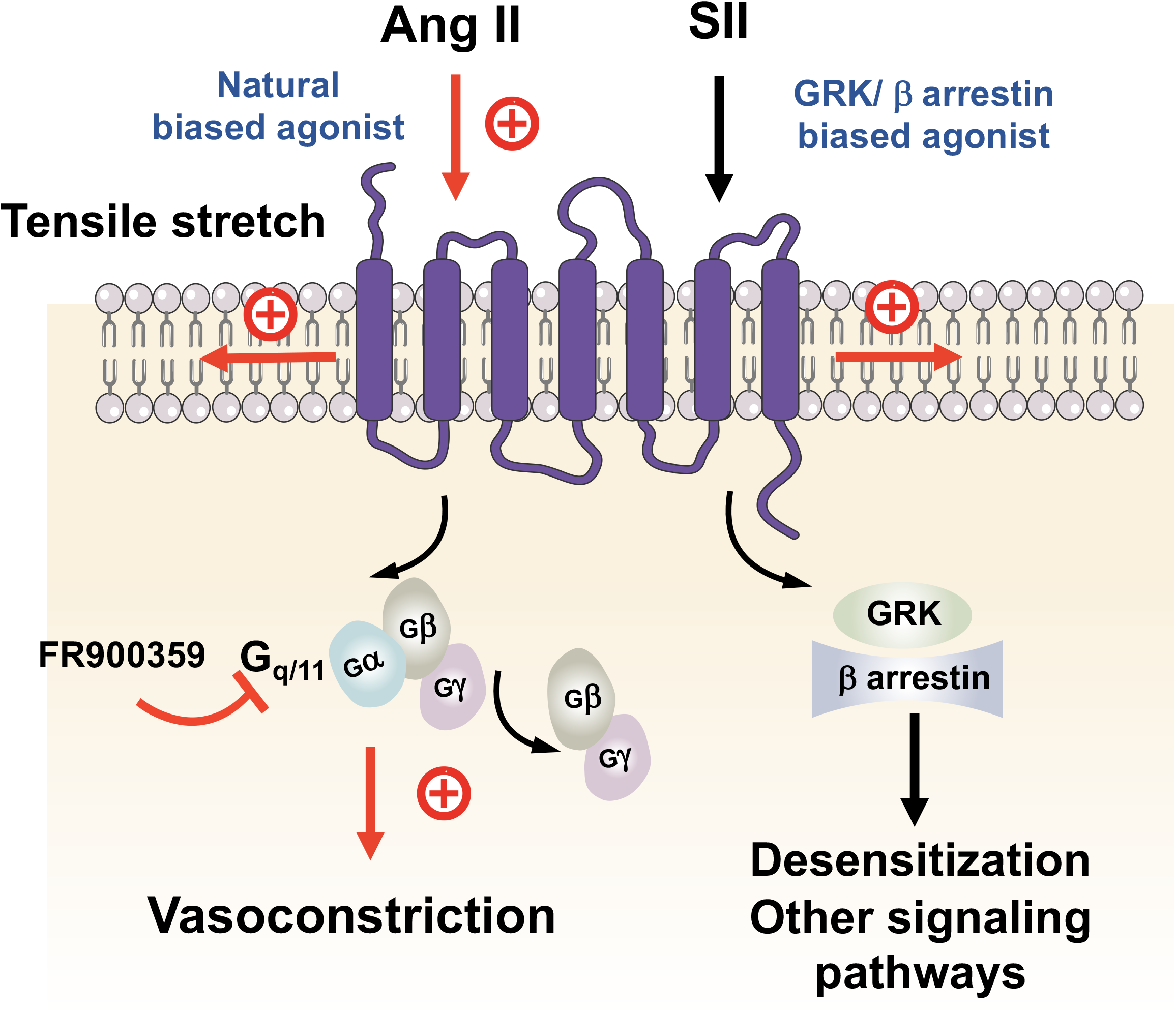
Schematic illustration of angiotensin II type 1a receptor (AT1aR) biased signaling cascade regulating myogenic arterial tone. Canonical G_q/11_ signaling pathway of the AT1R (purple blue) causes myogenic vasoconstriction whereas noncanonical β-arrestin-biased signaling is not involved in this process. G_q/11_ proteins are heterotrimeric G proteins, which are made up of alpha (α), beta (β) and gamma (γ) subunits. The alpha subunit is attached to either a guanosine triphosphate (GTP) or guanosine diphosphate (GDP), which serves as an on-off switch for the activation of the G-protein. Upon activation of the AT1aR by either ligand-independent mechanical stretch or the natural-biased ligand angiotensin II (Ang II), the Gβy complex is released from the Gα subunit after its GDP-GTP exchange for canonical G protein signaling to cause myogenic and/or humoral (Ang II-mediated) vasoconstriction. This pathway is inhibited by the G_q/11_ inhibitor FR900359. Although, GRKs and arrestins play a role in multiple noncanonical signaling pathways in cells, this pathway is unlikely engaged by mechanoactivated AT1Rs in response to tensile stretch or their natural ligand angiotensin II to cause vasoconstriction.

We hypothesized that G_q/11_ signaling contributes to myogenic tone in mesenteric and renal arteries and consistent with this idea, we found that the vasoconstrictor responses were strongly increased by the G_q/11_ AT1R biased agonists TRV120055 and TRV20056 (**Figure 10**). Moreover, we found that the G_q/11_ blocker FR900359 inhibited both myogenic tone and Ang II induced constrictions in mesenteric arteries and renal arterioles (**Figure 10**). The data imply that myogenic vasoconstriction requires canonical G_q/11_ signaling of the AT1aR. Consistently, myogenic tone is increased in the absence of regulator of G-protein signaling 2 (RGS2), which is an endogenous terminator of Galpha_q/11_ (Gα_q/11_) signaling (19) (37). The data align with findings indicating that mechanically activated AT1R generate diacylglycerol, which in turn activates protein kinase C (PKC) and induces the actin cytoskeleton reorganization necessary for pressure-induced vasoconstriction (22). Finally, our conclusions are supported by findings indicating that another G_q/11_-protein inhibitor YM 254890 profoundly reduced myogenic tone in mesenteric arteries (49). Note, this data contrast with recent findings, which proposed that G_12/13_- and Rho/Rho kinase-mediated signaling is required in myogenic vasoconstriction by inhibition of myosin phosphatase (5). The reason for the discrepancy is presently unknown, but may depend on which vessel order was utilized, i.e. 3^rd^ or 4^th^ order mesenteric versus 1^st^ or 2^nd^ order mesenteric arteries. Moreover, the myogenic response was only reduced by 50% in G_12/13_-deficient cerebral arteries (5), which may indicate that this pathway may play a role in some but not all vessels. Thus, it is possible that the relevance to the two signaling pathway differs between various vascular beds and artery branches. Our study provides firm evidence that AT1aRs coupled to G_q/11_ signaling is an essential component of dynamic mechanochemical signaling in arterial vascular smooth muscle cells causing myogenic tone (**Figure 10**).

Signaling of most GPCRs *via* G proteins is terminated (desensitization) by the phosphorylation of active receptor by specific kinases (GPCR kinases, or GRKs) and subsequent binding of ß-arrestins that selectively recognize active phosphorylated receptors. Although, GRKs and ß-arrestins play also a role in multiple noncanonical signaling pathways in the cell, both GPCR-initiated and receptor-independent (32) (15), our study failed to demonstrate that this pathway plays an important role in the myogenic response (**Figure 10**). Thus, it is unlikely that blood pressure lowering effects of β-arrestin biased AT1R agonists, e.g. Trevena 120027 (4), are caused by direct effects of this GPCR in the arterial smooth muscle cells.

In summary, we provide new and firm evidence for a mechanosensitive function of AT1aR in myogenic vasoconstriction in mesenteric, renal and cerebral arteries, i.e. in three different highly myogenic vascular beds. Our study clearly shows that mechanical stress activates AT1R in arterial smooth muscle cells, which subsequently triggers canonical G_q/11_ signaling, irrespective of GRK/β-arrestin signaling, to cause myogenic vasoconstriction. Our results argue against the idea of multiple mechanosensors coupled to noncanonical β-arrestin pathways generating myogenic arterial tone. These findings lay ground for additional studies to characterize the molecular mechanisms of mechanoactivated AT1aR coupled to G_q/11_ signaling in intact arteries, which may reveal new molecular targets for drug development to alleviate increased or dysregulated arterial tone in hypertension and other cardiovascular diseases.

## Acknowledgments

The Deutsche Forschungsgemeinschaft (DFG) supported our study (M.G.). We thank Thomas Coffman for providing *Agtr1a*^-/-^, *Agtr1b*^+/+^ and *Agtr1a*^flox^ mice. We thank Gabriele M. König and Evi Kostenis for FR900359.

